# Human Decidual RUNX1 Promotes Angiogenesis and Trophoblast Differentiation by Regulating Extracellular Vesicle Signaling

**DOI:** 10.1101/2025.07.09.663949

**Authors:** Jacob R. Beal, Xiangning Song, Athilakshmi Kannan, Indrani C. Bagchi, Milan K. Bagchi

## Abstract

During early pregnancy, human endometrial stromal cells differentiate into secretory decidual cells via a process regulated by ovarian steroid hormones. Decidual cells play a crucial role by secreting various factors that support essential events in forming a functional placenta, including uterine angiogenesis and the differentiation and development of trophoblasts. We previously reported that the conditional ablation of the transcription factor RUNX1 in the mouse uterus leads to subfertility due to insufficient maternal angiogenesis and impaired trophoblast differentiation. In this study, we examined the role of RUNX1 in facilitating communication mechanisms among human decidual cells and other cell types present in the pregnant uterus. We demonstrated that RUNX1 regulates the conserved HIF2α-RAB27B pathway in primary human endometrial stromal cells (HESC) during decidualization, which promotes the secretion of extracellular vesicles (EVs) by these cells. Consequently, the depletion of RUNX1 in HESC led to reduced EV secretion. Mass spectrometry identified several cargo proteins in decidual EVs, including ANGPTL2 and IGF2, which could regulate angiogenesis or trophoblast differentiation. We found that RUNX1 directly regulates their expression, resulting in partial changes to these cargoes when it is absent. We observed that delivering EVs lacking ANGPTL2 or IGF2 to human endothelial cells significantly decreased the formation of vascular networks compared to introducing control EVs carrying these factors. Furthermore, adding IGF2-depleted EVs to human trophoblast cells inhibited their differentiation into the extravillous trophoblast lineage. These findings collectively highlight the crucial role of decidual RUNX1 in promoting essential cell-cell interactions for angiogenesis and trophoblast differentiation during placenta formation.

## 1. INTRODUCTION

The uterine endometrium plays a major role during early pregnancy. In response to ovarian steroid hormones, estrogen and progesterone, the endometrium undergoes a remarkable transformation into the secretory decidua in a process known as decidualization. By relaying secreted factors to the fetus and other nearby maternal tissues, the decidua orchestrates critical processes within the pregnant uterus that are necessary for reproductive success. These processes include embryonic implantation, where the embryo attaches to the endometrial luminal epithelium and begins to invade deeper into the decidua (1,2), as well as maternal angiogenesis, which is essential for providing an increased blood supply to the uterus to meet the demands of the growing fetus (3,4). Both events are crucial for the development of the placenta, the transient organ responsible for supporting fetal growth throughout gestation.

Additionally, for proper placentation, the decidua acts in a paracrine manner to regulate the differentiation of the trophectoderm into appropriate subtypes (1,2). In humans, trophoblast stem (TS) cells can differentiate into two main lineages: the multinucleated syncytiotrophoblast (ST) and the invasive extravillous trophoblast (EVT). STs play a crucial role in the exchange of nutrients and gases between maternal and fetal blood (7), while EVTs are vital for their capability to invade deeper into maternal tissue, remodeling spiral arteries to redirect maternal blood flow toward the fetus (8).

Our understanding of the decidual factors that help coordinate the complex process of placentation is constantly evolving. Uterine-specific conditional knockout (cKO) mouse models have revealed several proteins whose expression in the decidua is crucial for proper placentation. These include ALK5 (3), Bmal1 (4), Nodal (5), Rac1 (6), and BMPR2 (7), among others. A recent report from our lab revealed that the conditional ablation of runt-related transcription factor 1 (RUNX1) in the uterus, through progesterone receptor (PR)-mediated Cre expression, also resulted in abnormal placentation and severe sub-fertility observed in the cKO female mice. This seemed to result from two main phenotypes—deficient maternal angiogenesis and impaired trophoblast differentiation and invasion. (8).

Recent studies have highlighted the significance of extracellular vesicle (EV) signaling from the endometrium during pregnancy, particularly as a mechanism for initiating and supporting placentation by the maternal decidua. (9). EV signaling involves the transport of molecules like miRNA or proteins via extracellular vesicles (EVs) from a donor cell to a recipient cell, triggering a functional change in the recipient. Endometrial EVs have been shown to play a significant role in guiding maternal neoangiogenesis (15,16) and facilitating trophoblast differentiation and invasion during placentation (15,17), although the exact mechanisms remain unclear.

In mice and humans, there exists a conserved regulatory pathway for EV secretion in endometrial stromal cells (15,18). Specifically, the transcription factor hypoxia-inducible factor 2 alpha (HIF2α) is expressed under the hypoxic conditions of early pregnancy and enhances the expression of the protein RAB27B, which is one of the proteins responsible for transporting the multivesicular body from the cytosol to the cell membrane, facilitating the release of EVs into the extracellular space (15,18). This conserved pathway enables the endometrium to coordinate events during early pregnancy, such as maternal angiogenesis and trophoblast differentiation invasion.

In this study, we tested the hypothesis that endometrial stromal RUNX1 plays a similar role in coordinating maternal angiogenesis and trophoblast differentiation in a human in vitro model as it does in mice. We identified how RUNX1 coordinates these functions by establishing a relationship between RUNX1 and EV secretion, as well as protein cargo composition. Additionally, we utilized human in vitro culture systems to examine the mechanisms through which RUNX1-controlled EV secretion promotes angiogenesis and influences trophoblast differentiation.

## 2. METHODS

### 2.1 HESC Culture and Differentiation

Deidentified primary HESCs were collected in accordance with the guidelines for the protection of human subjects participating in clinical research, as approved by the institutional review board of Wake Forest School of Medicine. Samples were acquired from fertile women aged 28 to 42 years, with a parity of 1 to 2, during the proliferative stage of the menstrual cycle via biopsy. Donors provided written informed consent and were confirmed to show no signs of endometrial pathologies. HESCs were isolated as described previously (10,11). Cells from at least two different donors were utilized in most experiments.

HESCs were passaged under normoxic conditions (20% O_2_) and cultured in Dulbecco’s modified Eagle medium (DMEM)/F-12 (Gibco) supplemented with 5% charcoal dextran-stripped FBS (Atlanta Biologicals), 50 μg/mL penicillin and streptomycin (Invitrogen). To induce decidualization HESCs were treated with a differentiation cocktail consisting of 1 μM progesterone (Sigma-Aldrich), 10 nM 17-β-estradiol (Sigma-Aldrich), and 0.5 mM 8-bromo-adenosine-3’,5’-cyclic monophosphate (Sigma-Aldrich) in DMEM/F-12 medium (Gibco) supplemented with 2% Exosome-depleted FBS (Gibco) and cultured in a hypoxia incubator (3% O_2_) for 72 hours before sample collection. Media and treatments were replenished every 48 hours.

### 2.2 siRNA-mediated Knockdown

Primary HESCs were transfected by siRNA targeting RUNX1, ANGPTL2, or IGF2 gene transcripts or a scrambled siRNA control (Dharmacon) following the manufacturer’s protocol (siLentFect; Bio-Rad). In short, a final concentration of 20 nM siRNA mixed with siLentFect lipid reagent to transfect cells. After incubation overnight with siRNA-lipid mixture the media was replaced with media containing differentiation cocktail as described above.

### 2.3 RNA Isolation and qPCR Analysis

After 72 hours of differentiation, RNA was extracted from HESCS using TRIzol (Invitrogen) according to the manufacturer’s protocol. AffinityScript Mulitple Temperature Reverse Transcriptase kit (Agilent) was used following the manufacturer’s instructions to convert RNA to cDNA. Quantitative PCR analysis was performed on the cDNA using Power Sybr Green PCR master mix (Applied Biosystems) and gene-specific primers (IDT) with 36B4 being used as a housekeeping gene. For each treatment condition the mean cycle threshold (Ct) was calculated from three replicates of each sample. ΔCt was calculated as the mean Ct of the gene of interest subtracted by the mean Ct of the housekeeping gene. ΔΔCt was calculated as the difference of ΔCt of experimental and control groups. Fold change of gene expression relative to control was computed as 2^-ΔΔCt^. The data was then displayed as mean fold induction and standard error of the mean (SEM) calculated from 2-6 independent experiments.

### 2.4 EV Isolation, Quantitation, and Protein Cargo Identification

As described previously (11), conditioned media was collected from HESCs after 72 hours of differentiation. Conditioned media centrifuged at 3,000 g for 10 min and 16,500 g for 20 min at 4 °C. Following centrifugation, supernatant was used to extract EVs using the miRCURY Exosome Cell/Urine/CSF Kit (Qiagen) as described in the manufacturer’s protocol. Before liquid chromatography/mass spectrometry (LC/MS) analysis, harvested EVs were further purified by ultracentrifugation at 120,000 g for 90 min at 4 °C.

EV quantitation was performed using microfluidic resistive pulse sensing (MRPS) on a Spectradyne nCS1^TM^ instrument (Spectradyne LLC, CA, USA) according to the manufacturer’s instructions. EV pellets were resuspended in filtered PBS and further diluted with PBS supplemented with 1% Tween20. EV suspension was loaded on polydimethylsiloxane cartridges which were factory calibrated to quantify particles only within specific size ranges. We utilized the C-400 cartridge to quantify particles from 65-400 nm in size as previously described (12,13). Data are presented as average particle concentrations (p/mL) of repeated measurements of independent samples.

After EV isolation and purification, EV pellets were submitted to the Mass Spectrometry Laboratory at the University of Illinois at Urbana-Champaign. LC/MS proteomics data was analyzed by Mascot (Matrix Science) to identify EV cargo proteins.

### 2.5 Chromatin Immunoprecipitation

ChIP assays were performed on HESC collected after 72 hours of differentiation using the ab500 ChIP Kit (Abcam) following the manufacturer’s protocol. For immunoprecipitation, fragmented DNA was incubated overnight with 5 μg of anti-RUNX1 (Abcam Cat# ab272456, RRID:AB_3675503) or Rabbit IgG (Abcam Cat# ab172730, RRID:AB_2687931). Real-time PCR was performed after immunoprecipitation and DNA purification steps. Primers were designed spanning putative RUNX1 response elements within the regulatory region of the IGF2 gene and a positive control within the MIP1α regulatory region (14). Resulting signals were normalized to input DNA, and data were displayed as % input and SEM calculated from 3 independent experiments.

### 2.6 Endothelial Tube Formation Assay

To assess capillary-like tube formation ability, 33,750 human umbilical vein endothelial cells (HUVECs) were seeded onto Matrigel basement membrane (Corning) which had solidified for 30 min at 37 °C. Cells were incubated for 6 hours in presence of EVs (∼2×10^8^ particles) from either HESCs treated with RUNX1, IGF2, ANGPTL2, or control siRNA. We used Calcein-AM (Thermo-Fisher) to stain the cells. Random microscopic images were captured and, to compare the tube formation ability across treatment groups, the number of junctions, segments, and branches were calculated in ImageJ as described previously (15). Representative images are shown, and quantitation data is from 3-5 biological replicates.

### 2.7 Trophoblast Culture and Differentiation

Human trophoblast stem cells were acquired from Dr. Michael Soares’ laboratory at the University of Kansas Medical Center with the permission of Dr. Hiroaki Okae of Tohoku University who developed the line (16,17). Cells were cultured on culture dishes coated with 5 μg/mL mouse collagen IV (Corning). Stem cells remain in a highly proliferative state when cultured in complete medium consisting of DMEM/F-12, 0.2% FBS, 0.3% fatty acid–free bovine serum albumin (BSA) (Thermo Fisher), 1% ITS-X (Thermo Fisher), 0.8 mM valproic acid (Sigma), 50 ng/mL recombinant hEGF (Sigma), 1.5 mg/mL L-ascorbic acid (Sigma), 5 µM Y27632 (Reprocell), 0.1 mM 2-mercaptoethanol (Sigma), 1 µM SB431542 (Reprocell), 2 µM CHIR99021 (Reprocell), 0.5 µM A83-01 (Reprocell), and penicillin–streptomycin.

To induce trophoblast stem cells differentiation into EVT cells, cells were seeded in chamber slides (Ibidi) with polymer coverslips coated with 1 μg/mL collagen IV and grown on days 1 and 2 of differentiation in a differentiation media containing DMEM/F-12, 0.3% BSA, 1% ITS-X, 0.1 mM 2-mercaptoethanol, 2.5 µM Y27632, 7.5 µM A83-01, 4% KnockOut Serum Replacement (Thermo Fisher), 100 ng/mL hNRG-1(Cell Signaling Technology), and penicillin– streptomycin supplemented with 2% Matrigel (Corning). On day 3, the media was replaced with the same media, excluding NRG1, and supplemented with 0.5% Matrigel. On Day 6, media was replaced with the same media, excluding NRG1 and KnockOut Serum Replacement, and supplemented with 0.5% Matrigel. Media throughout the differentiation process was supplemented with EVs (∼2×10^8^ particles) derived from HESCs treated with RUNX1, IGF2, or control siRNA. On day 8, cells were analyzed for markers of EVT differentiation.

EVTs were fixed with 3.7% formaldehyde and permeabilized with 0.3% Triton-X 100 in PBS for 5 min, followed by blocking with PBS + 2% FBS. Cells were incubated overnight with antibodies against HLA-G (Novus Biologicals; RRID: NBP1-43123), MMP2 (Cell Signaling Technology; RRID: D4M2N) or Integrin αV (Cell Signaling Technology; RRID:4711). Nuclei were stained with DAPI and slides were mounted by non-hardening mounting medium. Imaging was performed on Zeiss 700 confocal microscope and relative fluorescent quantitation was done in ImageJ.

### 2.8 Statistical Analysis

Statistical analysis was performed in all experiments using an unpaired Student’s t-test to determine if treatments were statistically different from the control. GraphPad Prism Version 9 (GraphPad Inc.) was utilized for statistical analysis. A P-value of ≤ 0.05 was needed to be considered statistically significant.

## 3. RESULTS

### 3.1 RUNX1 acts upstream of HIF2α/RAB27B to control EV secretion by HESCs

In RUNX1 cKO mice, ablation of RUNX1 from endometrial stromal cells induced abnormal responses from other cell types within the uterus. Namely, mesometrial endothelial cells exhibited deficient vascular network formation and trophoblast cells displayed reduced invasion into the uterine tissue (8). This phenomenon suggests the possibility that the effects of endometrial stromal RUNX1 may be related to paracrine signaling, which would explain how other tissue’s function is impaired by the absence of endometrial stromal RUNX1.

For this reason, we first investigated an important cell signaling mechanism that is conserved in both mice and human endometrial stromal cells during decidualization, EV secretion. Within decidualizing endometrial stromal cells, transcription factor HIF2α is known to regulate the expression of *RAB27B* gene, which codes for a vesicular trafficking protein crucial to the secretion of EVs (12,13). We aimed to determine if this EV secretion pathway is dysregulated by the absence of RUNX1 in decidualizing HESCs. To do this we used a well-established system to induce decidualization in HESCs. A differentiation cocktail (DC) consisting of 1 μM progesterone, 10 nM 17-β-estradiol, and 0.5 mM 8-bromo-adenosine-3’,5’-cyclic monophosphate was added to HESC media, and after 72 hours of differentiation relevant gene transcripts were quantified.

Before induction of the decidualization program, HESCs were treated with 20 nM of scrambled control siRNA or siRNA specifically targeted to *RUNX1* transcripts. Treatment with 20 nM *RUNX1* siRNA led to a robust ∼80% knockdown of *RUNX1* gene expression as quantified by qPCR (**Fig. 1A**). Additionally, we found that *RUNX1* knockdown caused a reduction in *HIF2α* expression (**Fig. 1B**), which led to a concomitant decrease in *RAB27B* gene expression (**Fig. 1C**). After 72 hours of differentiation, we also collected the conditioned media from the cells.

**Fig. 1.**
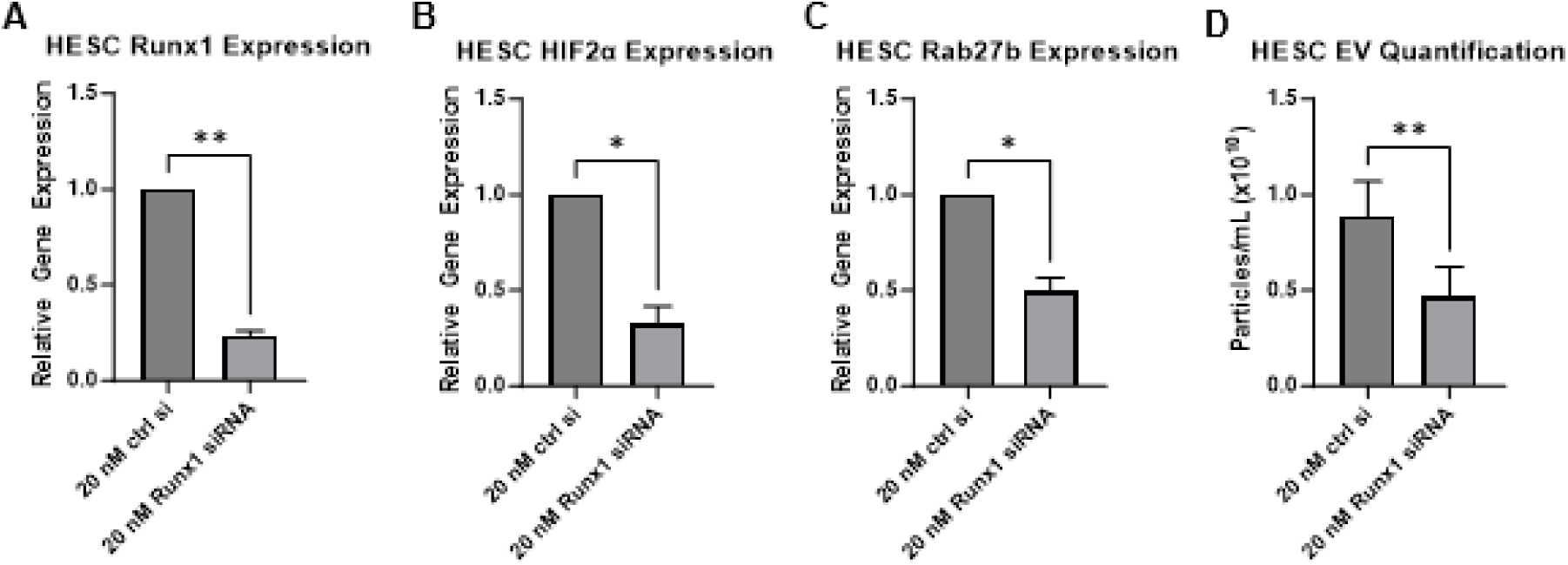
RUNX1 mediates EV secretion in differentiating HESCs by regulating *HIF2α*/*RAB27B* gene expression. HESCs were treated with 20 nM control or *RUNX1*-specific siRNA for 24 h, then grown in culture under hypoxic conditions with Decidualization Cocktail for 72 h. RNA was extracted and gene expression analysis was performed using primers specific for *RUNX1* (**A**), and members of an EV secretion pathway in differentiating HESCs, transcription factor *HIF2α* (**B**), and vesicular trafficking protein *RAB27B* (**C**). EVs were also isolated from conditioned media of *RUNX1*-specific or control siRNA using the miRCURY (Qiagen) kit and analyzed by MRPS (Spectradyne, LLC). Concentration of EVs ranging from 60 to 400 nm was quantified (**D**). Data shown as mean fold change ± SEM (n = 3-6 independent experiments), * p<0.05, ** p<0.01 relative to control siRNA treatment.

We isolated EVs from the conditioned media using a precipitation-based kit (miRCURY) and, after resuspension in PBS, quantified the EV concentration through microfluidic resistive pulse sensing (MRPS). We found that treatment with *RUNX1* siRNA caused decidualizing HESCs to secrete about half as many EVs as control siRNA-treated HESCs (**Fig. 1D**). Together, these data suggest an important mechanism by which human decidual RUNX1 may promote uterine angiogenesis and trophoblast differentiation, regulating the secretion of EVs.

### 3.2 Absence of RUNX1 in HESC alters specific EV protein cargoes

EVs contain a variety of cargoes, including proteins, RNA, and lipids, which can induce a functional change in the recipient cells. EVs isolated from endometrial cells have been shown to contain many proteins which are known to play roles in key processes during early pregnancy, such as angiogenesis, trophoblast differentiation, extracellular matrix remodeling, and decidualization (13,18). Alteration to the EV protein cargo profile can result in dysfunction in the processes typically guided by endometrial EV secretion, potentially leading to adverse reproductive outcomes (9,19).

Therefore, we aimed to determine if, in addition to the regulation of EV secretion, RUNX1 also plays a role in determining the protein cargoes of EVs secreted by decidualizing HESCs. After treatment with *RUNX1* or control siRNA, followed by the addition of DC, as described above, conditioned media were collected from the treated HESCs. We proceeded to isolate EVs from the conditioned media using a precipitation-based kit. After isolation, we further purified the EVs by subjecting them to ultracentrifugation. The resultant EV pellet was then analyzed by liquid chromatography-mass spectrometry (LC-MS) to identify the protein cargoes of the EVs.

Comparison of protein cargoes isolated from EVs secreted by cells treated with control siRNA, deposited at (20) or *RUNX1* siRNA, deposited at (20), revealed a number of differential protein cargoes between the two treatment groups. 875 proteins were found to be common EV cargoes regardless of the expression level of RUNX1 in the HESCs, while 199 cargoes were unique to EVs from control siRNA-treated cells, and 298 cargoes were unique to EVs derived from HESCs lacking RUNX1 (**Fig. 2A** and deposited at (20)).

**Fig. 2.**
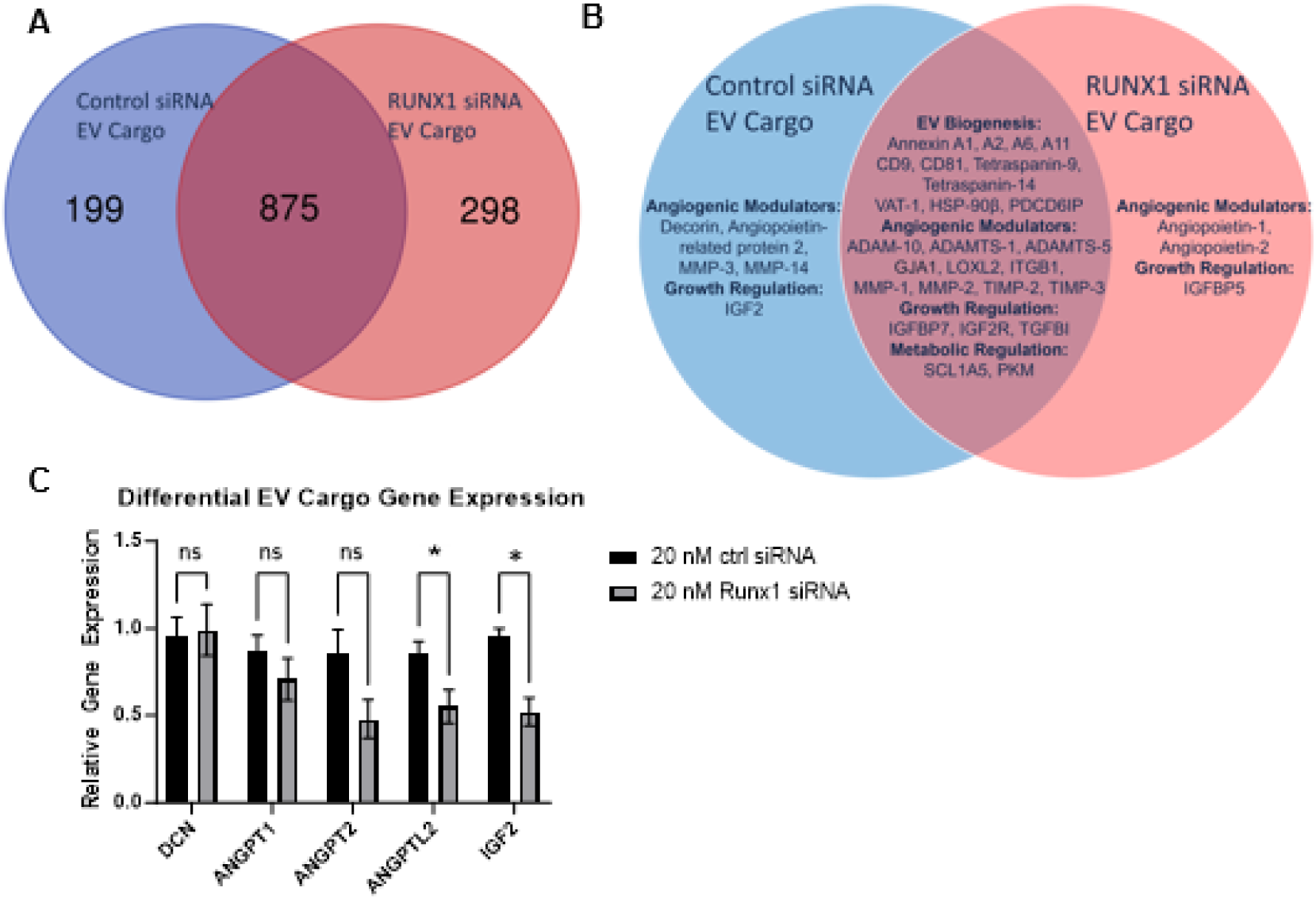
RUNX1 partially influences HESC EV protein cargo composition via gene expression regulation. HESCs were treated with 20 nM control or *RUNX1*-specific siRNA for 24 h then grown in culture under hypoxic conditions with Decidualization Cocktail for 72 h. EVs were isolated from conditioned media of *Runx1*-specific or control siRNA using ultracentrifugation. Proteomic analysis identified cargo proteins present in either or both samples (**A**). A partial list of potentially functional protein cargoes is provided (**B**). RNA was extracted and gene expression analysis was performed using primers specific for genes encoding select differential EV cargoes, *ANGPT1 and ANGPT2* (**C**), as well as *DCN*, *ANGPTL2*, and *IGF2* (**D**). *36B4* was used to normalize gene expression. Data shown as mean fold change ± SEM (n = 3-6 independent experiments), * p<0.05, relative to control siRNA treatment.

Among the shared EV cargo proteins, we found several proteins involved with EV biogenesis, including various annexins, tetraspanins, as well as classical markers of EVs, CD9 and CD81. Also included in the shared proteins were angiogenic modulators, metabolic regulators, and growth factors and binding proteins; all of which have been previously identified as EV cargoes in decidualizing HESCs (13) and appear to be unaffected by the absence of RUNX1 (**Fig. 2B, center**).

We also found that, while some proteins were present as EV cargoes in both control and *RUNX1* siRNA-treated groups, there were some changes to the amount of each protein detected within EVs after *RUNX1* siRNA treatment. Interestingly, when examining the fold change in the EV cargo load of proteins known to be angiogenic modulators (**Table 1**), we found that several proteins traditionally associated with promotion of angiogenesis, Angiopoietin-related protein 2 (ANGPTL2), Decorin (DCN), Insulin-like growth factor II (IGF2), and Matrix metalloproteinases 3 and 14 (MMPs), were found to be absent after *RUNX1* siRNA treatment, while traditional negative regulators of angiogenesis, Metalloproteinase inhibitor 2 and 3 (TIMPs) were found to be increased in EVs after siRNA treatment. However, surprisingly, there were a couple of positive promoters of angiogenesis, Angiopoietin 1 and 2 (ANGPTs), that were found in HESC EVs after *RUNX1* siRNA treatment. Similarly, when examining the fold change in EV cargo load of proteins known to be influencers of trophoblast differentiation (**Table 2**), we found that Erbb2 receptor tyrosine kinase 4 (ERBB4) and IGF2, factors vital for trophoblast differentiation were absent in HESC EVs after *RUNX1* siRNA treatment. In contrast, inhibitors of IGF2 signaling, cation-independent mannose-6-phosphate receptor (IGF2R) and Insulin-like growth factor-binding protein 5 (IGFBP5) were increased in EVs after *RUNX1* siRNA treatment.

**Table 1.** RUNX1 induces changes in EV cargo load of potentially angiogenic proteins in differentiating HESCs. A partial list of EV cargoes with potential for angiogenic function secreted by *RUNX1*-specific siRNA HESCs compared to control siRNA treated HESCs as identified by mass spectrometry after 72 hours of differentiation. Fold change is shown as the number protein matches in EVs isolated from *RUNX1*-specific siRNA HESCs divided by number of protein matches in EVs isolated from control siRNA HESCs. Inf refers to proteins that were only present as HESC EV cargo after treatment with RUNX1-specific siRNA.

| Gene Name | Protein Description | Fold Change |
| --- | --- | --- |
| **Angiogenic Modulators** |  |  |
| ANGPT1 | Angiopoietin-1 | inf |
| ANGPT2 | Angiopoietin-2 | inf |
| TIMP2 | Metalloproteinase inhibitor 2 | 4 |
| TIMP3 | Metalloproteinase inhibitor 3 | 2.5 |
| LOXL2 | Lysyl oxidase homolog 2 | 1.928571429 |
| MMP2 | 72 kDa type IV collagenase | 1.454545455 |
| ADAMTS1 | A disintegrin and metalloproteinase with thrombospondin motifs 1 | 1.428571429 |
| ITGB1 | Integrin beta | 1.333333333 |
| MMP1 | Interstitial collagenase | 1.333333333 |
| ADAM10 | Disintegrin and metalloproteinase domain-containing protein 10 | 1 |
| GJA1 | Gap junction alpha-1 protein | 1 |
| ADAMTS5 | A disintegrin and metalloproteinase with thrombospondin motifs 5 | 0.25 |
| ANGPTL2 | Angiopoietin-related protein 2 | 0 |
| DCN | Decorin | 0 |
| IGF2 | Insulin-like growth factor II | 0 |
| MMP3 | Stromelysin-1 | 0 |
| MMP14 | Matrix metalloproteinase-14 | 0 |

**Table 2.** RUNX1 induces changes in EV cargo load of proteins with potential to modulate trophoblast differentiation in differentiating HESCs. A partial list of EV cargoes with potential to modulate trophoblast differentiation secreted by *RUNX1*-specific siRNA HESCs compared to control siRNA treated HESCs as identified by mass spectrometry after 72 hours of differentiation. Fold change is shown as the number protein matches in EVs isolated from *RUNX1*-specific siRNA HESCs divided by number of protein matches in EVs isolated from control siRNA HESCs. Inf refers to proteins that were only present as HESC EV cargo after treatment with RUNX1-specific siRNA.

| Gene Name | Protein Description | Fold Change |
| --- | --- | --- |
| Regulators of Trophoblast Differentiation |  |  |
| IGFBP5 | Insulin-like growth factor-binding protein 5 | inf |
| IGF2R | Cation-independent mannose-6-phosphate receptor | 3.5 |
| IGFBP7 | Insulin-like growth factor-binding protein 7 | 1.5 |
| CDC42 | Cell division control protein 42 homolog | 1.333333333 |
| TGFBI | Transforming growth factor-beta-induced protein ig-h3 | 1 |
| EIF2S3 | Eukaryotic translation initiation factor 2 subunit 3 | 1 |
| EIF4A1 | Eukaryotic initiation factor 4A-I | 1 |
| ERBB4 | Erb-b2 receptor tyrosine kinase 4 | 0 |
| IGF2 | Insulin-like growth factor II | 0 |

We specifically further investigated angiogenic modulators ANGPTL2 and DCN, as well as growth factor IGF2, which may also play a role in trophoblast differentiation in addition to angiogenesis. These proteins were found in the control-treated EVs but were no longer present in the EV cargoes from *RUNX1* siRNA-treated HESCs (**Fig. 2B, left**). This implies that the presence of RUNX1 is required for these particular proteins to be included in the HESC EV cargoes. We hypothesized that this might be due to RUNX1, as a transcription factor, regulating the gene expression of proteins. As a result of the ablation of RUNX1, there would be a decrease in the cytoplasmic accumulation of proteins whose expression is regulated by RUNX1. This would make it less likely that these proteins are available to be enclosed within the multivesicular body and eventually secreted within EVs.

To test this hypothesis, we analyzed the gene expression of several relevant proteins that were found to be differentially present in the EV cargoes of our two treatment groups. Surprisingly, we had identified two angiogenic modulators, Angiopoietin-1 and Angiopoietin-2, that were only present in EVs from *RUNX1* siRNA-treated HESCs (**Fig. 2B, right**). However, upon further investigation, we found that absence of RUNX1 did not induce an increase in gene expression of these factors (**Fig. 2C**). Similarly, the gene expression of the angiogenic modulator *DCN* was unchanged after *RUNX1* siRNA treatment, despite DCN only being present in EVs isolated from control treated HESCs (**Fig. 2D**). Interestingly, the other two relevant factors who had been identified to be present as EV cargoes in control treated HESCs but absent in EVs from HESCs treated with *RUNX1* siRNA, ANGPTL2 and IGF2, both had decreased gene expression in the absence of RUNX1 (**Fig. 2D**). This suggests that during decidualization, RUNX1 normally promotes *ANGPTL2* and *IGF2* gene expression. ANGPTL2 is a key angiogenic factor (21), and IGF2, in addition to being an angiogenic factor, is a crucial regulator of trophoblast differentiation (22,23).

Together, these data suggest that the absence of RUNX1 induces changes to the EV protein cargo profile of HESCs. Specific proteins, such as ANGPTL2 and IGF2, are now absent from the HESC EV cargo due to RUNX1 no longer being present to promote their gene expression. As both proteins are likely to play roles in angiogenesis and trophoblast differentiation, this could be a cause of the phenotypes displayed in the previously reported RUNX1 cKO mouse model (8).

### 3.3 RUNX1 expression in HESCs enhances pro-angiogenic activity of HESC-EVs

Endometrial stromal regulation of uterine angiogenesis is crucial during early pregnancy. The development of an extensive vascular architecture within the uterus is necessary to meet the demands of the growing embryo. Maternal vasculature is essential for supporting the embryo prior to placentation and serves as a maternal blood supply during the placentation process (24,25). The endometrial expression of vascular endothelial growth factor A (VEGF-A), gap junction A1 (GJA1), and various matrix metalloproteinases (MMPs) as well as fibroblast growth factors (FGFs) has been associated with the regulation of angiogenesis in the uterus during early pregnancy. (24,26–28).

We already observed that RUNX1 promotes the gene expression of angiogenic factors that are secreted from HESCs as EV protein cargoes. To determine if RUNX1 also contributes to the expression of angiogenic factors within the endometrium that may be released by non-EV secretion pathways, we performed siRNA-mediated knockdown of *RUNX1*. We observed a decrease in the gene expression of angiogenic factors VEGF-A, MMP2, and GJA1 (**Fig. 3A**), indicating that RUNX1 typically plays a crucial role in promoting their expression.

**Fig. 3.**
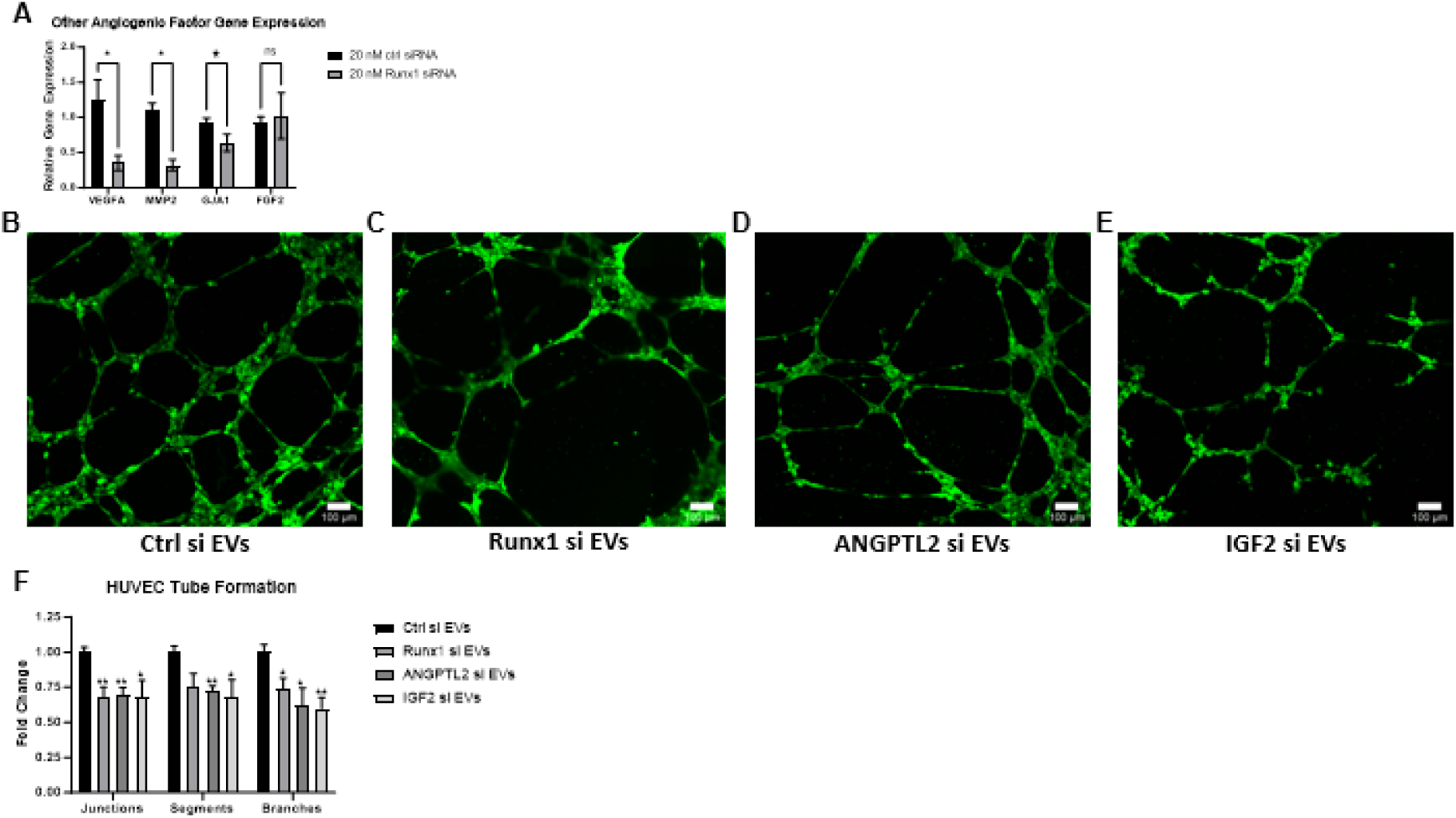
RUNX1 expression in HESCs enhances HESC-EVs pro-angiogenic activity by promoting the expression of angiogenic factors. HESCs were treated with 20 nM control or *RUNX1*-specific siRNA for 24 h then grown in culture under hypoxic conditions with Decidualization Cocktail for 72 h. **A.** RNA was extracted and gene expression analysis was performed using primers specific for known angiogenic factors that had not been previously identified to be differential protein cargoes, *VEGFA*, *MMP2*, *GJA1*, and *FGF2*. *36B4* was used to normalize gene expression. Data shown as mean fold change ± SEM (n = 3-6 independent experiments), * p<0.05, relative to control siRNA treatment. **B.** EVs were isolated from conditioned media of HESCs treated with control, *RUNX1*-, *ANGPTL2*-, or *IGF2*-specific siRNA using the miRCURY (Qiagen) kit. EVs were cultured at 3 x 10^11^ p/mL with HUVECs plated with basement membrane matrix. Representative images were taken of tube formation. **C**. Tube formation capability was quantified by counting the number of junctions, nodes, branches, and segments in ImageJ from 4 random microscopic images per replicate. Data shown as mean fold change ± SEM, * p<0.05, ** p<0.01.

In addition to regulating the gene expression of angiogenic factors in HESCs, we have shown that depletion of RUNX1 induces a change to the EV cargo protein profile, including the exclusion of important angiogenic factors ANGPTL2 and IGF2 from the cargo proteins. To assess whether the angiogenic promotion capability of HESC EVs is compromised due to the altered protein cargoes resulting from HESC RUNX1 depletion, we conducted a functional assay. Primary human umbilical vein endothelial cells (HUVECs) gradually form capillary-like structures when seeded onto a matrix membrane, and the addition of HESC-derived EVs enhances the tube formation process (13). We found that the addition of EVs derived from HESCs depleted of RUNX1 displayed impaired tube formation compared to the addition of an equivalent number of control EVs (**Fig. 3B-C**), as assessed by quantitation of junctions, segments, and branches formed by the HUVECs (**Fig. 3F**). Also, EVs derived from HESCs depleted of ANGPTL2 (**Fig. 3D, 5F**) or IGF2 (**Fig. 3E-F**) when added to the HUVEC culture exhibited reduced tube formation compared to the addition of control EVs.

These data confirm that RUNX1 promotes the expression of angiogenic factors in HESCs. This study identifies specific angiogenic factors, such as ANGPTL2 and IGF2, as critical RUNX1-regulated cargoes carried by EVs secreted from the HESCs. The presence of these angiogenic factors is crucial for HESC EVs’ ability to enhance angiogenesis, as demonstrated by our in vitro angiogenesis model.

### 3.4 RUNX1 regulates IGF2 cargo in EVs secreted by HESCs to modulate trophoblast differentiation into the EVT lineage

Throughout placentation, the maternal blood supply remains a source of nutrition for the growing embryo. In addition to the extensive vasculature architecture developed within the decidua, the fetal trophoblast cells must also invade deep into the decidua to remodel maternal spiral arteries and redirect blood flow towards the fetus. Specialized trophoblast cells called EVTs are primarily responsible for this invasion and remodeling (29). EVT differentiation from trophoblast progenitor cells is a tightly controlled process, and our lab has previously described that maternal-derived EVs markedly enhance the differentiation of trophoblast stem cells into EVTs (13).

The differentiation of trophoblast stem cells to EVT is characterized by an increased expression of IGF2 by EVTs, and autocrine secretion of IGF2 has long been thought to be a driver of human trophoblast invasiveness (30). Our findings suggest an interesting role for the contribution of maternal IGF2, packaged within HESC EVs, on EVT differentiation. We hypothesized that RUNX1-dependent expression and secretion of HESC IGF2 could influence EVT differentiation via EV-mediated cell-cell communication.

*IGF2* gene expression is regulated by RUNX1 in decidualizing HESCs (**Fig. 2D**). To determine if RUNX1 control of *IGF2* expression is a direct effect, we tested several putative RUNX1 binding sites in the *IGF2* gene regulatory region and found through chromatin immunoprecipitation (ChIP) studies that RUNX1 significantly occupies DNA within the *IGF2* regulatory region (**Fig. 4A**). These data suggest that RUNX1 is a direct regulator of *IGF2* gene expression in decidualizing HESCs.

**Fig. 4.**
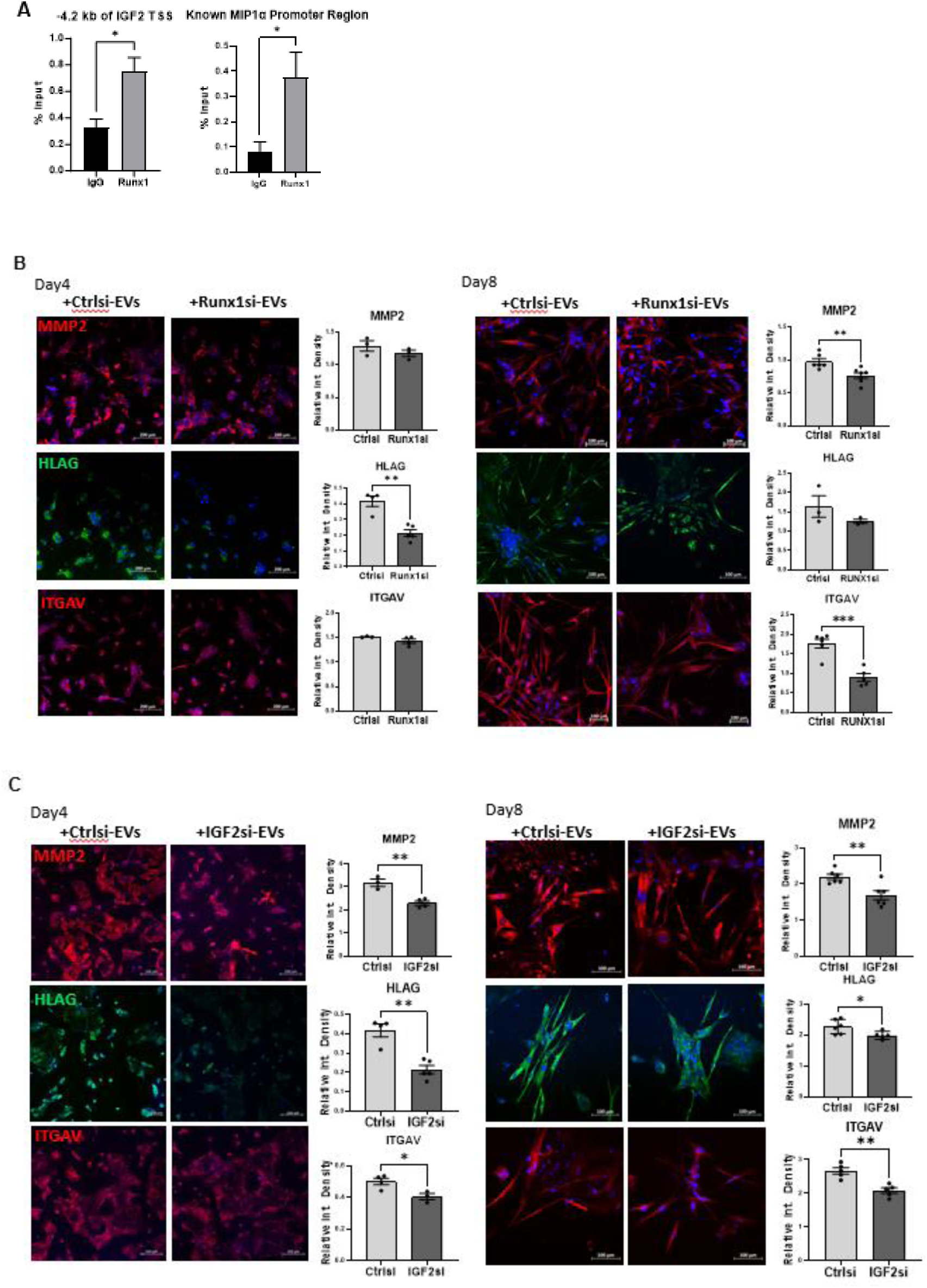
RUNX1 expression in HESCs modulates trophoblast differentiation into EVT lineage by regulating IGF2 expression. **A.** ChIP was performed on HESCs that had been treated with Decidualization Cocktail for 72 h. Chromatin enrichment was quantified by qPCR. Primers flanking RUNX1 consensus sequences within the IGF2 promoter or the positive control MIP1α regulatory region were used to quantify chromatin enrichment. Representative data shown as % input ± SEM (n = 3 independent experiments), * p<0.05, relative to IgG. **B-C.** Primary trophoblast stem cells were induced to differentiate into EVTs and were incubated with EVs isolated from HESCs treated with control, *RUNX1*-specific siRNA (**B**), or *IGF2*-specific siRNA (**C**). Expression of EVT markers MMP2, HLAG, and ITGAV was quantified by immunocytochemistry at the halfway point (Day 4) and completion (Day 8) of the differentiation process. The relative fluorescence was quantified by ImageJ and normalized to DAPI. Data is shown as mean fold change ± SEM relative to treatment with control siRNA EVs, **p<0.01, ***p<0.001.

To test whether the expression of RUNX1 in HESCs altered the ability of HESC EVs to influence trophoblast function, we utilized a primary human trophoblast stem cell line derived by Drs. Hiroaki Okae and Takahiro Arima. These cells are maintained in a self-renewing stem state and can be differentiated into EVTs by an 8-day differentiation process (16).

Over the course of differentiation, cells exhibit morphological changes as well as an increase in expression of markers of EVT differentiation, *HLAG*, *MMP2*, and *ITGAV*. We compared the expression of these markers in response to supplementation with EVs derived from control, *RUNX1* siRNA-treated (**Fig. 4B**), or *IGF2* siRNA-treated (**Fig. 4C**) HESCs. As shown in **Fig. 4B**, EVs from *RUNX1* siRNA-treated HESCs were moderately less efficient in promoting EVT differentiation compared to control EVs. At the midpoint of differentiation (Day 4), only the *HLAG* marker was differentially expressed between the treatments; however, by the end of differentiation (Day 8), this difference was not apparent. In contrast, the control EVs appear to be efficiently promoting the expression of both *MMP2* and *ITGAV* markers at later stages of differentiation compared to those secreted by *RUNX1* siRNA-treated HESCs.

Remarkably, as seen in **Fig. 4C**, EVs derived from *IGF2* siRNA-treated HESCs displayed a more prominent deficiency in enhancing EVT differentiation. Throughout the differentiation process, all markers of EVT differentiation were reduced in the cells treated with EVs derived from *IGF2* siRNA-treated HESCs compared to control EVs, suggesting a significant role for IGF2 as a maternal EV cargo in promoting trophoblast to EVT differentiation. Collectively, these data point to a pathway where endometrial stromal RUNX1-regulated expression and secretion of IGF2 as EV cargo enhances the differentiation of trophoblast stem cells to the EVT lineage during early pregnancy.

## 4. DISCUSSION

Endometrial stromal cells play a central role during early pregnancy, secreting a variety of proteins that serve as mediators of various processes essential for reproductive success. As such, it is crucial to understand the function of proteins within endometrial stromal cells, as well as to determine how their role can impact the function of other cell types within the uterus. Valuable insights into these interactions can be made from *in vivo* rodent studies; however, not all processes are conserved between species, and validation in a human *in vitro* system remains an important step in furthering our understanding of the cellular mechanisms that operate during early pregnancy in humans.

Here, we built upon the exciting recent findings of Kannan et al. (8), who reported that conditional knockout of transcription factor RUNX1 within the murine uterus led to severe subfertility due to deficient maternal vascularization and impaired trophoblast differentiation and development. Using a well-established human *in vitro* model of endometrial decidualization, we discovered that the role of decidual RUNX1 is indeed conserved between mice and humans, with downregulation of *RUNX1* expression in decidualizing HESCs leading to a muted capacity of HESCs to promote angiogenesis and differentiation in human endothelial and trophoblast cells, respectively. Additionally, we were able to provide unique insights into the mechanism by which HESC RUNX1 coordinates these processes.

A major finding of this study is that RUNX1 plays a role in regulating EV secretion in HESCs. We found that, in decidualizing HESCs, RUNX1 acts upstream of the HIF2α/RAB27B pathway that is conserved in decidual cells of both mice and humans to promote EV secretion. This finding enhances our understanding of how ablation of decidual RUNX1 had such profound consequences to maternal angiogenesis and trophoblast differentiation and development in the RUNX1 cKO mouse model described by Kannan et al. (8). EV secretion is a crucial method of cell-to-cell communication during early pregnancy and its dysregulation can lead to dire consequences to the health of a pregnancy (31). To our knowledge, we are the first to report the involvement of RUNX1 in endometrial stromal cell EV secretion; however, mutations in the RUNX1 gene have been previously linked to defects in dense granule secretion in platelets (32). Interestingly, platelet dense granule secretion is also regulated by RAB27B (33), providing another example of RUNX1 promoting RAB27B-controlled secretion pathways.

In addition to a reduced quantity of EVs secreted into the extracellular space, we also found that downregulation of RUNX1 transcripts led to an altered EV cargo profile. Specific modulators of angiogenesis, such as ANGPTL2 and IGF2, were no longer found as EV cargoes in HESCs that had been treated with RUNX1 siRNA. ANGPTL2 has been previously reported to promote tube formation in endothelial cells (34,35). IGF2 has also been linked to the growth of vasculature (22) and it has been shown to promote the transition of trophoblast cells into an invasive lineage (30), making it a crucial protein that likely controls multiple processes during early pregnancy. We found that the gene expression of *ANGPTL2* and *IGF2* is directly related to *RUNX1* expression. However, a few other differential EV protein cargoes that we identified had gene expression that was unaffected by RUNX1, suggesting that RUNX1 is only *indirectly* involved in these proteins’ inclusion as EV cargoes through another unidentified mechanism.

One of the most important aspects of the changing EV protein cargo profile that we have observed in HESCs treated with *RUNX1* siRNA is that these alterations to EV cargoes lead to a muted response in the recipient cells. Using an *in vitro* tube formation assay, we showed that EVs derived from HESCs treated with *RUNX1* siRNA failed to promote tube formation to the same extent as control HESC EVs. In various biological contexts, RUNX1 has been reported to promote angiogenesis (13,40–42). Consistent with these previous reports, we found that RUNX1 regulated the gene expression of several angiogenic modulators. However, in this study, we have discovered that part of RUNX1’s ability to promote angiogenesis in HESCs is mediated by EV signaling. Not all angiogenic modulators are generally contained within HESC EVs, but two significant angiogenic modulators, ANGPTL2 and IGF2, were found in control HESC EVs, but not EVs derived from HESCs treated with *RUNX1* siRNA. After further investigation, we found that EVs from ANGPTL2- or IGF2-depleted HESCs also failed to promote tube formation by HUVECs to the same extent as control EVs. These data suggest that RUNX1 facilitates uterine angiogenesis by promoting the expression of various angiogenic factors in HESCs. The inclusion of at least two of these factors, ANGPTL2 and IGF2, as EV cargoes is essential for the maximal capacity of HESC EVs to promote uterine angiogenesis.

HESC EVs are also known to enhance the differentiation of human trophoblast stem cells (13) Consistent with the phenotypic defects observed by Kannan et al. in mice lacking uterine Runx1 (13), we found that RUNX1 depletion in HESCs resulted in a reduced enhancement of human trophoblast differentiation into EVTs by HESC EVs. Notably, we found that while treating trophoblast stem cells with EVs derived from HESCs treated with RUNX1 siRNA somewhat diminished the enhancement of trophoblast differentiation by HESC EVs, EVs from IGF2-depleted HESCs dramatically reduced trophoblast stem cell differentiation into the EVT lineage. From these data, we postulate that the majority of RUNX1’s influence on trophoblast differentiation comes directly from its regulation of IGF2 expression and secretion. Incomplete knockdown of *IGF2* gene expression in HESCs, which occurs after treatment with *RUNX1* siRNA, appears to be sufficient to partially mute the influence of HESC EVs on the EVT differentiation process. However, a more complete knockdown of *IGF2* gene expression occurs after HESC treatment with *IGF2* siRNA, and that directly corresponds to a more dramatic reduction in the ability of HESC EVs to promote EVT differentiation. Traditionally, it has been suggested that only autocrine secretion of IGFs by the embryo is responsible for driving trophoblast proliferation and differentiation (36–38). However, our discovery that RUNX1-regulated IGF2 inclusion as an EV cargo is crucial for HESC EV enhancement of EVT differentiation suggests that the role of maternal paracrine IGF2 signaling in trophoblast differentiation may be more significant than previously believed.

In conclusion, our study has revealed an important role for human decidual RUNX1 in regulating essential processes during early pregnancy, demonstrating conservation of decidual RUNX1’s profound impact on angiogenesis, trophoblast differentiation, and development that was previously uncovered in murine studies. We have also further elucidated the molecular mechanisms by which RUNX1 influences these processes; namely, RUNX1’s regulation of EV secretion and protein cargo composition, particularly the inclusion of key factors ANGPTL2 and IGF2 as EV cargoes. We have also provided intriguing evidence highlighting the significant contribution of maternal IGF2 to trophoblast differentiation, challenging existing paradigms and inviting future investigation. Overall, this study has established that human decidual RUNX1 is a key factor during early pregnancy, influencing uterine angiogenesis and trophoblast differentiation through the regulation of EV-mediated communication. This research contributes valuable insights to our growing understanding of the regulation of EV-mediated communication, which is necessary for orchestrating the complex series of events crucial for successful pregnancy.

